# Functional expression of opioid receptors and other human GPCRs in yeast engineered to produce human sterols

**DOI:** 10.1101/2021.05.12.443385

**Authors:** Björn D.M. Bean, Colleen J. Mulvihill, Riddhiman K. Garge, Daniel R. Boutz, Olivier Rousseau, Brendan M. Floyd, William Cheney, Elizabeth C. Gardner, Andrew D. Ellington, Edward M. Marcotte, Jimmy D. Gollihar, Malcolm Whiteway, Vincent J.J. Martin

**Affiliations:** Department of Biology, Centre for Applied Synthetic Biology, Concordia University, Montréal, QC, H4B1R6, Canada; Department of Molecular Biosciences, Center for Systems and Synthetic Biology, The University of Texas at Austin, Austin, Texas, 78712, USA; DEVCOM Army Research Laboratory-South, Austin, 78712, TX, USA; Center for Molecular and Translational Human Infectious Diseases Research, Department of Pathology and Genomic Medicine, Houston Methodist Research Institute, Houston Methodist Hospital, Houston, TX, USA

**Keywords:** humanized yeast, opioid biosensor, G protein-coupled receptors, sterol pathway engineering

## Abstract

The yeast *Saccharomyces cerevisiae* is a powerful tool for studying G protein-coupled receptors (GPCRs) as they can be functionally coupled to its pheromone response pathway. However, some exogenous GPCRs, including the mu opioid receptor, are non-functional in yeast, which may be due to the presence of the fungal sterol ergosterol instead of the animal sterol cholesterol. We engineered yeast to produce cholesterol and introduced the human mu opioid receptor, creating an opioid biosensor capable of detecting the peptide DAMGO at an EC_50_ of 62 nM and the opiate morphine at an EC_50_ of 882 nM. Furthermore, introducing mu, delta, and kappa opioid receptors from diverse vertebrates consistently yielded active opioid biosensors that both recapitulated expected agonist binding profiles with EC_50_s as low as 2.5 nM and were inhibited by the antagonist naltrexone. Additionally, clinically relevant human mu opioid receptor alleles, or variants with terminal mutations, resulted in biosensors that largely displayed the expected changes in activity. We also tested mu opioid receptor-based biosensors with systematically adjusted biosynthetic intermediates of cholesterol, enabling us to relate sterol profiles with biosensor sensitivity. Finally, cholesterol-producing and sterol intermediate biosensor backgrounds were applied to other human GPCRs, resulting in SSTR5, 5-HTR4, FPR1 and NPY1R signaling with varying degrees of cholesterol dependence. Our sterol-optimized platform will be a valuable tool in generating human GPCR-based biosensors, aiding in ongoing receptor deorphanization efforts, and providing a framework for high-throughput screening of receptors and effectors.

## INTRODUCTION

G-protein coupled receptors (GPCRs) detect diverse extracellular stimuli, modulating signal transduction pathways that allow cells to respond to their environment. These seven transmembrane domain proteins typically function by binding external ligands, which induce conformational changes, propagating a signal across the plasma membrane and triggering internal signaling pathways^1^. Owing to their critical functions and their ubiquity as the largest family of human membrane proteins, one third of current FDA approved therapeutic targets are GPCRs^2^. Yet, while these targets are functionally understood, discovery of new GPCR-interacting therapeutics remains challenging in part due to screening limitations.

Assays of GPCR activity in the yeast *S. cerevisiae* may accelerate the search for therapeutics by allowing simple, cheap, and high-throughput screens. Commonly, these assays are based on functionally linking GPCRs to the yeast pheromone response pathway (PRP). Normally in the PRP, a native GPCR binds a mating pheromone, causing a GTP-GDP substitution in the Gα protein Gpa1, triggering a mitogen activated protein kinase signaling cascade culminating in upregulation of Ste 12-regulated genes^3,4^. This pathway can be commandeered to make a biosensor by replacing the native GPCR, creating a chimeric Gpa1 to maintain the GPCR-G protein interaction, and placing a reporter under the control of a Ste 12-regulated promoter^5^. Such yeast-based biosensor designs, initially applied to the β2-adrenergic receptor^6^, have now been used for over 50 receptors^7^. Yet, in many cases GPCRs cannot be functionally expressed in yeast^8–10^. This may be due to poor expression or folding^11–13^, defects in trafficking to the plasma membrane^10^ or differences in the chemical environment^14–16^.

In particular, the function of heterologously-expressed human GPCRs may be disrupted by the membrane lipid composition in yeast, as the dominant sterol is ergosterol, as opposed to cholesterol^9^. This would be consistent with past work documenting the importance of cholesterol-GPCR interactions^17–19^ and the frequent presence of cholesterol molecules as elements of established GPCR structures^20^. Thus, modifying the sterol profile of yeast may increase the proportion of human GPCRs that can be functionally expressed. Previously, deletion of the ergosterol biosynthetic genes *ERG5/6*, and introduction of zebrafish enzymes DHCR7/24 resulted in yeast producing cholesterol up to 96% of total sterol content^21,22^. While this modification disrupted the function of the endogenous GPCR Ste2^21^, its effect on heterologous GPCRs has not been tested.

It would be valuable to apply yeast-based rapid screening approaches to human opioid receptors. The main opioid receptor types, mu, delta, and kappa, are all GPCRs implicated in nociception and analgesia^23^. Drugs targeting these receptors, and the human mu opioid receptor (HsMOR) in particular, are essential front-line pain treatment medicines, but have also enabled misuse and dependence^24^. Expansion of available drugs that target these receptors but lack the side-effects of prototypical opioids could help resolve these issues. Though an HsMOR-based biosensor would provide a powerful tool for identifying new drug candidates, past efforts to construct this tool have failed in part due to the sterol composition of yeast membranes leading to low HsMOR activity^9^.

Here we describe a new biosensor background based on signaling through the PRP in a yeast strain engineered to produce cholesterol. This background dramatically improves HsMOR activity relative to an ergosterol-rich strain, enabling the characterization of structural and clinically-relevant HsMOR variants. We probed the agonist sensitivities of opioid biosensors based on 15 different receptors and found that opioid receptor activity and agonist specificities are well conserved in yeast. Screening a library of HsMOR-based biosensors with different sterol profiles allowed us to uncover how cholesterol intermediates affect signaling and establish that the cholesterol producing background was highly effective. Lastly, we applied the cholesterol-producing background as a platform to study other human GPCRs.

## RESULTS

### Construction of an opioid biosensor in a cholesterol-producing yeast

Previous work found that yeast-expressed human mu opioid receptor (HsMOR) was only active in lysates when ergosterol was removed and cholesterol added^9^. Therefore, we investigated whether HsMOR may be active in yeast cells engineered to produce cholesterol instead of ergosterol, and if active, whether linking HsMOR to the PRP would create an opioid biosensor.

We made a biosensor chassis based on previous studies linking GPCRs to the PRP^5,25^ (Figure 1A). The pheromone receptor, Ste2, was removed to avoid interference and the final five residues of the Gα protein, Gpa1, were swapped with the endogenous HsMOR-interacting protein G_iα3_ (K_468_IGII>ECGLY) to generate a chimera previously shown to link exogenous GPCRs to the PRP^5^. We chose green fluorescent protein (GFP) expression as an output and selected the promoter controlling expression by using alpha mating factor to screen the ability of eight highly PRP-regulated promoters to express GFP^26,27^ (Figure 1B). While p*FUS1* is often used^5,6,28^, we found that *pAGA1, pFIG1* and *pFIG2* all yield roughly four times the response, leading us to select *pFIG1::GFP* as the reporter.

**Figure 1.**
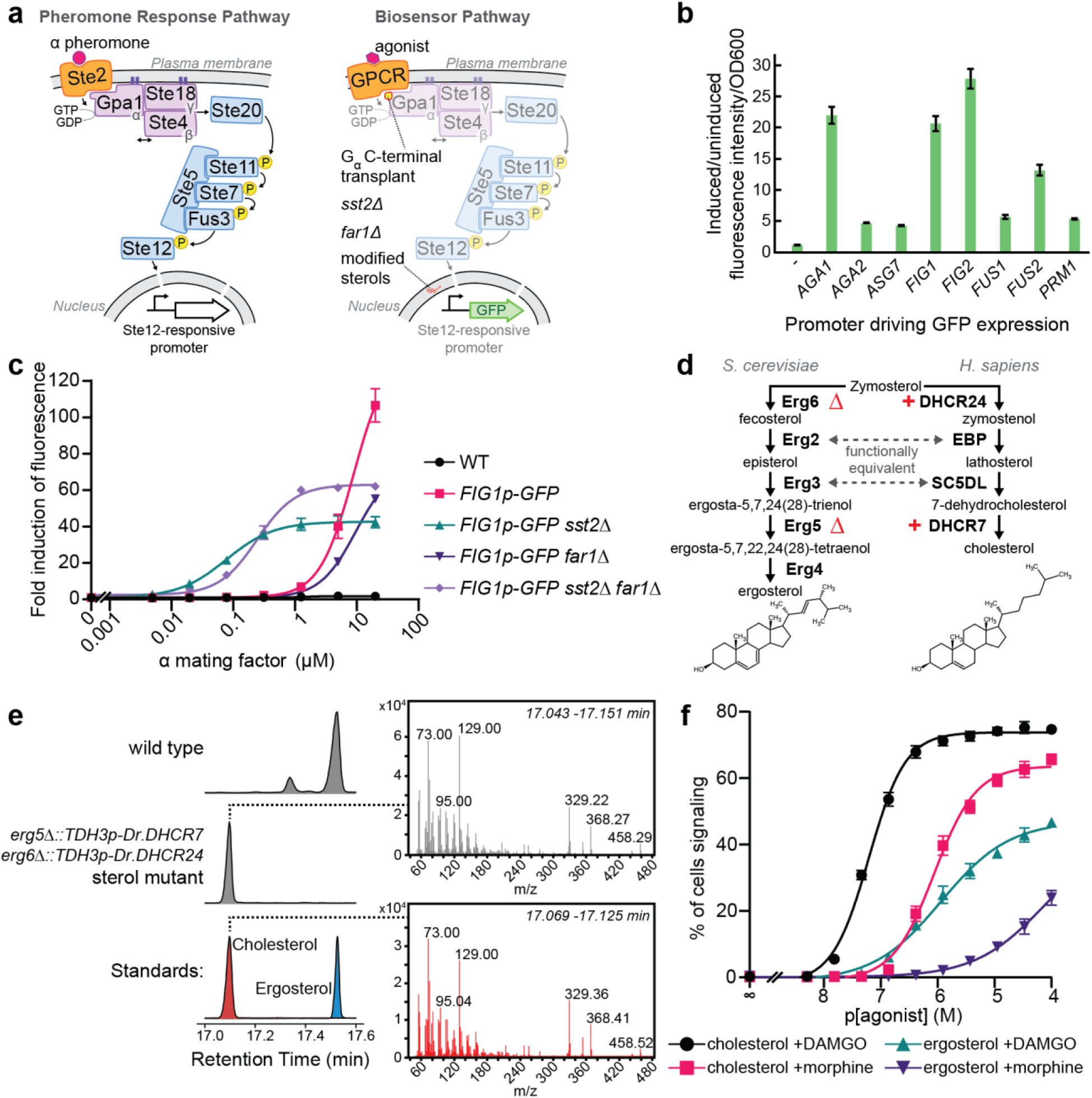
Development of a yeast-based opioid biosensor. **(a)** Strategy to adapt the yeast pheromone response pathway (PRP, left) into a biosensor pathway (right). An exogenous GPCR is introduced and linked to the pathway by a Gpa1 chimera. Deletion of *SST2* potentiates signaling and deletion of *FAR1* blocks signaling-induced cell cycle arrest. Ergosterol is replaced by cholesterol and a promoter controlled by the PRP-responsive Ste12 transcription factor is connected to a GFP output. **(b)** Biosensor reporter promoters tested using alpha mating factor-induced GFP expression from yeast Ste12-responsive promoters after 3h treatment. n=3. **(c)** PRP sensitivity and activity resulting from *SST2* and/or *FAR1* deletions. Strains were treated with alpha mating factor for 6 h, mean fluorescence was measured by flow cytometry. n=3; 10000 cells/strain/replicate. **(d)** Ergosterol and cholesterol biosynthetic pathways from zymosterol. **(e)** GC-MS analysis of sterol extracts showing successful synthesis of cholesterol in an *erg5/6 DHCR7/24* background. Chromatograms indicating retention times of derivatized sterols from a wild type strain (top), the engineered strain (middle), and standards (bottom). MS spectra extracted from engineered strain (top) and cholesterol standard (bottom). **(f)** DAMGO and morphine dose-response curves for ergosterol and cholesterol-producing HsMOR-based biosensors. Measured by flow cytometry after 8 h agonist treatment. n=3; 10000 cells/strain/replicate. Error bars indicate SEM.

The chassis was optimized by deleting *FAR1* and *SST2*, respectively preventing PRP-induced cell cycle arrest and increasing sensitivity by reducing pathway deactivation. While these deletions are a common strategy^7^ their effects on heterologous signaling are poorly documented, so we measured how they influenced *pFIG1::GFP* response to alpha mating factor (Figure 1C). As expected, *SST2* deletion increased sensitivity (17.9x) though background fluorescence also increased (4.2x), which limited fold induction of fluorescence. While Far1 is not prescribed a role in pheromone sensitivity we found that *FAR1* deletion decreased sensitivity both in wild type and *sst2* backgrounds (7.0x and 3.1x respectively). The background fluorescence of the *sst2* strain was also reduced by *FAR1* deletion from 4.2x to 1.6x that of wild type. This, together with the ability of a *far1* strain to facilitate longer assays, led us to select a *sst2 far1* background even though the *FAR1* deletion impacts sensitivity.

Next, the strain was engineered to produce cholesterol instead of ergosterol. Cholesterol and ergosterol are structurally similar, with zymosterol as the last common intermediate (Figure 1D). Following Souza et al., we deleted *ERG5/6* and added *TDH3p*-driven zebrafish DHCR7/24 to block ergosterol production and redirect zymosterol to cholesterol^22^. In this modified cholesterol production pathway Erg2 and Erg3 fulfill the roles of human EBP and SC5DL respectively. GC-MS analysis showed 94% of sterols were cholesterol with 4% dehydrolathosterol also present (Figure 1E; Supplementary Figure 4).

Addition of yeast codon-optimized *OPRM1*, the gene encoding HsMOR, driven by the strong *CCW12* promoter resulted in a candidate opioid biosensor. The sensitivities of both this cholesterol membrane biosensor, and a native yeast membrane (ergosterol) biosensor, were assessed by measuring fluorescence after 8 hours exposure to different concentrations of the HsMOR agonists [D-Ala^2^, N-MePhe^4^, Gly-ol]-enkephalin (DAMGO) and morphine. Importantly, initial tests indicated a strong pH dependence, with optimal morphine signaling at pH 7.1, as opposed to the normal yeast growth media pH of 5-5.5 (Supplementary Figure 1B). We postulate that improved biosensor signaling at a pH of 7.1 results from conditions that more closely resemble the conditions HsMOR is exposed to in the brain (pH 7.2 intracellular^29^, pH 7.4 extracellular^30^).

With pH adjustment, both ergosterol and cholesterol-producing biosensors responded to DAMGO and morphine (Figure 1F). Consistent with the known cholesterol dependence of HsMOR^9^, the cholesterol-rich biosensor was dramatically more effective, with a lower EC_50_ (62 ± 3 nM vs 1.3 ± 0.3 μM DAMGO; 0.9 ±0.1 μM vs ~110 ± 40 μM morphine) and a larger proportion of cells signaling. The presence of any signaling in the ergosterol strain was unexpected given that previously [^3^H] DAMGO binding by HsMOR had not been detected in yeast^9^. The absence of binding may have been due to the use of a higher buffer pH (7.5) and/or lower receptor expression. Taken together, we have constructed two opioid biosensors with different detection limits that demonstrate conversion of sterols to cholesterol can improve human GPCR function in yeast.

### An array of biosensors based on different opioid receptors reveals fidelity of agonist selectivity

Next we expanded the set of receptors being tested to explore the degree of opioid receptor functional conservation in yeast. Opioid receptors exist throughout Vertebrata; we selected a diverse group, including five of each type: mu (MOR), kappa (KOR) and delta (DOR). Within each type are receptors from humans (*Homo sapiens*, Hs), mice (*Mus musculus*, Mm), and zebrafish (*Danio rerio*, Dr). Additional receptors were included from the cow (*Bos taurus*, Bt), flying fox (*Pteropus vampyrus*, Pva), bearded dragon (*Pogona vitticeps*, Pvi), Burmese python (*Python bivittatus*, Pb), and Mexican tetra, (*Astyanax mexicanus*, Am). As expected from the high degree of opioid receptor conservation, a MUSCLE-generated^31^ phylogenetic tree showed segregation by receptor type (Figure 2A). Furthermore, MORs and DORs clustered closely, consistent with the current model of MORs and DORs emerging from a common ancestral receptor^32^.

**Figure 2.**
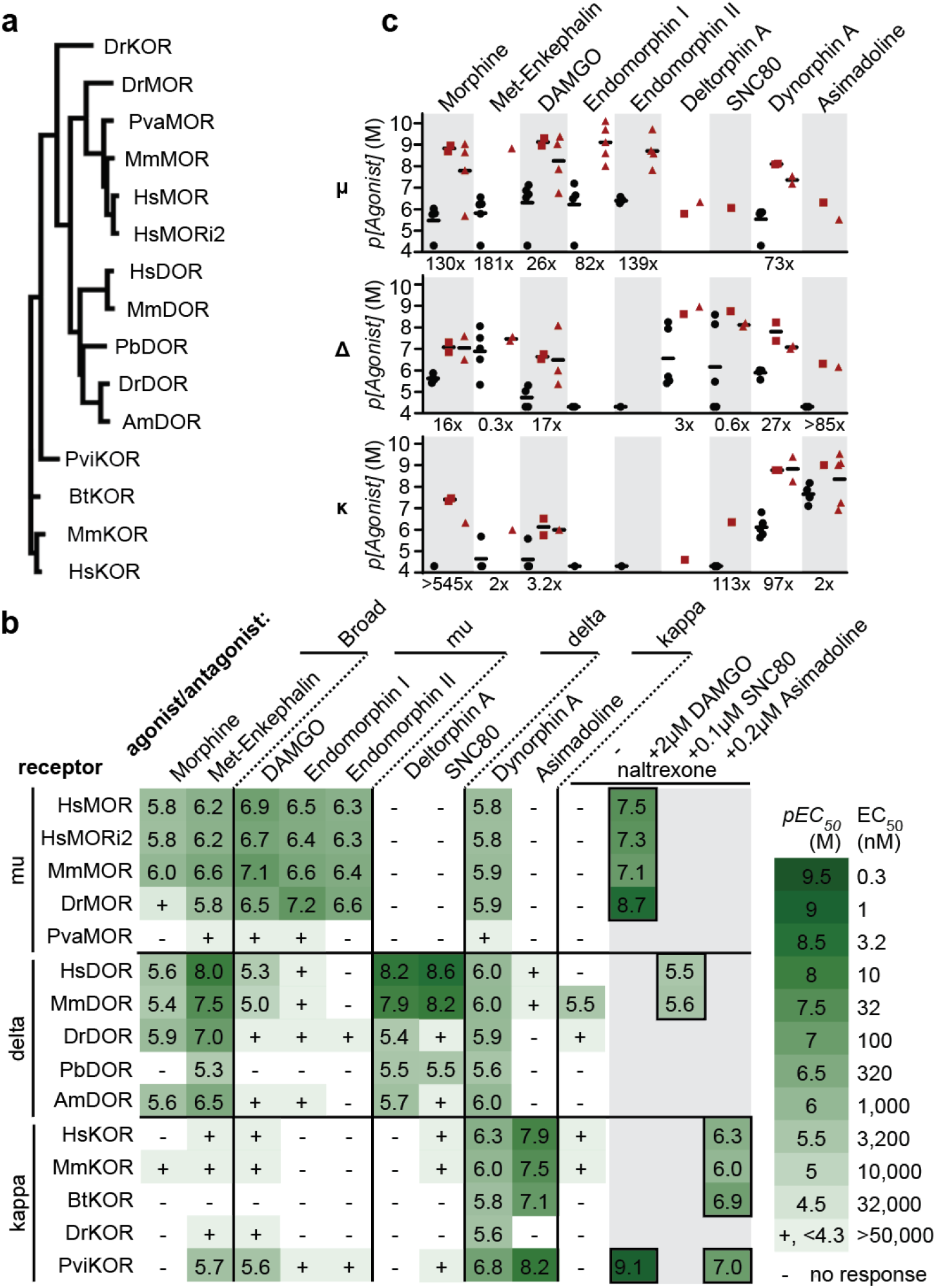
Activity of opioid receptors in the cholesterol-producing strain. **(a)** A protein-based phylogenetic tree of opioid receptors selected to make biosensors. MOR, mu receptor, KOR, kappa receptor, DOR, delta receptor. Dr, *D. rerio*, zebrafish; Pva, *P. vampyrus*, flying fox; Mm, *M. musculus*, mouse; Pb, *P. bivittatus*, python; Am, *A. mexicanus*, Mexican tetra; Pvi, *P. vitticeps*, bearded dragon; Bt, *B. taurus*, cow. i2, isoform 2. **(b)** Activity of agonists and the antagonist naltrexone on opioid receptors assayed by flow cytometry. Agonists clustered by literature receptor-type specificity. n=3, >2073 cells/condition/replicate. **(c)** Biosensor agonist sensitivities (black circles) relative to literature binding constants (red squares) and EC_50_/IC_50_ values (red triangles)^34–45^. Fold decrease in sensitivity of the best biosensor relative to the literature average is indicated.

Biosensors based on these opioid receptors were tested for activity and agonist specificity. Agonists were selected based on human receptor specificities: morphine and met-enkephalin are broad-acting^33^, DAMGO and Endomorphins I/II are MOR-specific^34,35^, Deltorphin A and SNC80 are DOR-specific^36,37^, and Dynorphin A and Asimadoline are KOR-specific^38,39^. Most were short 4-17 residue peptides, or peptide-based, except for the benzylisoquinoline alkaloid morphine and the two heterocycles SNC80 and Asimadoline. For each biosensor-agonist pair a dose-response curve was made and EC_50_ was calculated (Figure 2B, Supplementary Figure 1/2). Remarkably, all strains showed some response to at least three of the agonists tested, and, with the exception of the PvaMOR strain, all biosensors were sensitive enough to use agonist concentrations below 50 μM to determine EC_50_.

Agonist specificities largely matched those of human receptors in endogenous conditions, though receptor sensitivity was reduced (Figure 2B/C). MORs and DORs responded to the broad acting agonists morphine and met-enkephalin with EC_50_s of 30 nM to 3 μM, while KORs responded poorly, consistent with KORs’ reported low met-enkephalin sensitivity but not their reported 47-538 nM morphine sensitivity^34,35^. MOR-specific agonists were detected by MORs with 60-500 nM EC_50_s while other receptor types were less sensitive (EC_50_ >5 μM). Likewise, only DORs fully responded to the HsDOR-specific agonists deltorphin A and SNC80 (EC_50_s 2.5 nM - 4 μM). KOR-based biosensors were most sensitive to KOR-specific agonists with EC_50_s as low as 6.3 nM, though most biosensors responded to Dynorphin A, consistent with reported MOR and DOR Dynorphin A sensitivity^34^. While agonist specificities were maintained, biosensors displayed type-specific decreases in receptor sensitivity relative to values reported for more native environments, with DORs performing best (11x decrease) followed by KORs (43x decrease) and MORs (105x decrease)^34–45^ (Figure 2C).

The effects of an antagonist, naltrexone, were also determined^34^. Alone, naltrexone occasionally functioned as a partial agonist, at most eliciting a signaling population one seventh the size of that induced by a strong agonist (Figure 2B, Supplementary Figure 1E). Antagonist activity was tested by incubating biosensors for eight hours with an amount of agonist sufficient for strong signaling (2 μM DAMGO, 0.1 μM SNC80, 0.2 μM asimadoline) and varying concentrations of naltrexone (Figure 2B, Supplementary Figure 1E). Naltrexone blocked activity in all cases and, in line with binding coefficients previously determined in CHO cells^34^, MORs were most sensitive (IC_50_: 2 - 80 nM), followed by KORs (IC_50_: 0.8 - 500 nM) and DORs (IC_50_: 2.5 - 3.2 μM). Together, the ability of these biosensors to reconstitute both agonist specificities and antagonist activity make them powerful tools for assessing how opioid receptors in native environments will respond to a compound.

### Signal sequences disrupt mu opioid receptor function

Although the biosensors recapitulated the pattern of response seen in vertebrates, sensitivity was lower than in native cells, suggesting aspects of receptor expression or the signaling environment could be improved. To explore if opioid receptor sensitivity was limited by expression or localization defects, GFP-tagged HsMOR (HsMOR-GFP) was imaged in a cholesterol-producing background. HsMOR-GFP primarily localized to the ER with a secondary vacuolar pool (Figure 3D). The unexpected lack of HsMOR-GFP on the plasma membrane, where functional GPCRs have previously been observed^10^, suggests a folding or trafficking defect may be leading to endoplasmic reticulum (ER) retention and/or misdirection to the vacuole. GFP tagging itself is unlikely to be causing mislocalization as the tag only reduced biosensor response to DAMGO by 34% (Supplementary Figure 1C). Given the degree of HsMOR-GFP ER retention, we speculated that increasing plasma membrane localization might improve biosensor function.

**Figure 3.**
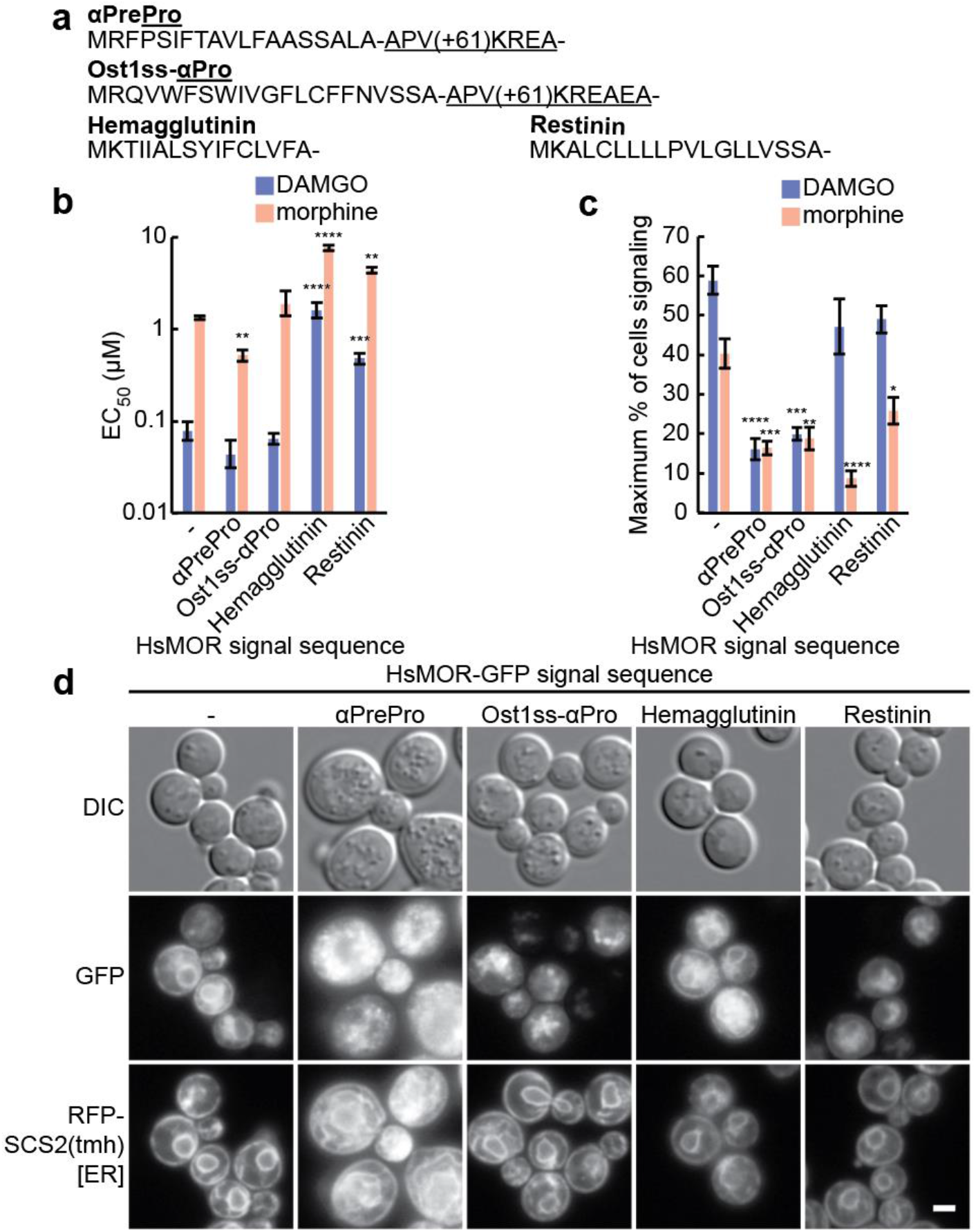
Effect of signal sequences on HsMOR function. **(a)** N-terminal signal sequences tested on HsMOR. **(b)** Effect of signal sequences on HsMOR biosensor sensitivity to DAMGO and morphine. Unpaired one-way ANOVA of *pEC_50_*s: n=3, >7595 cells/strain/replicate; P < 0.0001 for each agonist; Dunnett’s multiple comparisons tests against WT shown. **(c)** Effect of signal sequences on HsMOR signaling population. Unpaired one-way ANOVA: n=3, >7595 cells/strain/replicate; P < 0.0001 (DAMGO) and P < 0.001 (morphine); Dunnett’s multiple comparisons tests against WT shown. **(d)** Imaging of HsMOR-GFP with the indicated signal sequences in a cholesterol-rich background with the ER marker RFP-SCS2(tmh). Error bars indicate SEM. Scale bar is 2 μm. *, P < 0.05; **, P < 0.01; ***, P < 0.001; **** P < 0.0001.

GPCR expression and localization can be improved by appending N-terminal signal sequences^10,46^, short peptides that mediate ER insertion^47^. While integral membrane proteins, such as opioid receptors, often lack signal sequences because transmembrane helices are sufficient for ER targeting, adding the sequences can increase ER insertion speed, minimizing misfolding^47,48^. Therefore, the effects of appending signal sequences to HsMOR were assayed. We tested sequences from yeast (α-mating factor pre-pro, αPrePro; Ost1 signal peptide - α-mating factor pro, Ost1ss-αPro) as well as others previously used to improve GPCR expression in mammalian cells (influenza Hemagglutinin; Restinin)^46,49^ (Figure 3A).

Appending signal sequences to HsMOR generally did not improve sensitivity to either DAMGO or morphine, and was instead disruptive in two distinct ways (Figure 3B/C). Only αPrePro increased sensitivity, by roughly two-fold for both agonists, whereas Ost1ss-αPro was neutral and the Hemagglutinin and Restinin sequences caused 320-fold decreases in sensitivity. While the yeast αPrePro and Ost1ss-αPro sequences were neutral or beneficial for sensitivity, they dramatically decreased the maximum size of the signaling population, by 72% and 66% respectively. In contrast, the Hemagglutinin and Restinin sequences didn’t significantly affect the DAMGO-induced signaling population, selectively disrupting the morphine response. Taken together, the signal sequence classes have contrasting effects: yeast-based sequences had a neutral or positive effect on sensitivity and a reduced signaling population, whereas the Hemagglutinin and Restinin sequences disrupted sensitivity but did not always reduce the signaling population.

To better understand this dichotomy, signal sequence-tagged HsMOR-GFP was imaged in a cholesterol-producing background (Figure 3D). Strikingly, the αPrePro and Ost1ss-αPro sequences resulted in enlarged granular cells, expanded ER membranes and relocalization of HsMOR-GFP to puncta. In contrast, the Hemagglutinin and Restinin tags did not disrupt cellular morphology and HsMOR-GFP remained ER-localized, though the vacuolar pool may have increased. These results suggest that the yeast-based sequences cause global cellular disruptions, perhaps through partial HsMOR-GFP misfolding, which may be associated with premature ER exit. Cellular stress likely disrupts signaling, leading to the observed reductions in biosensor signaling competency. The other sequences did not disrupt cellular morphology and consequently did not display consistent decreases in the biosensor signaling population. The link between cellular localization and sensitivity was unclear. Overall, while the yeast signal sequences subtly improved HsMOR sensitivity, the associated cellular disruptions decreased the signaling population such that the signal sequences were not beneficial.

### Biosensors recapitulate the effects of missense mutants in HsMOR

Our biosensor platform may enable convenient characterization of rare opioid receptor alleles. Introduction of receptor variants should allow measurement of altered receptor sensitivities and signaling strength, potentially predicting clinically relevant changes in responses to analgesics. To probe our platform’s ability to detect these changes we tested known HsMOR missense mutations (Figure 4A).

**Figure 4.**
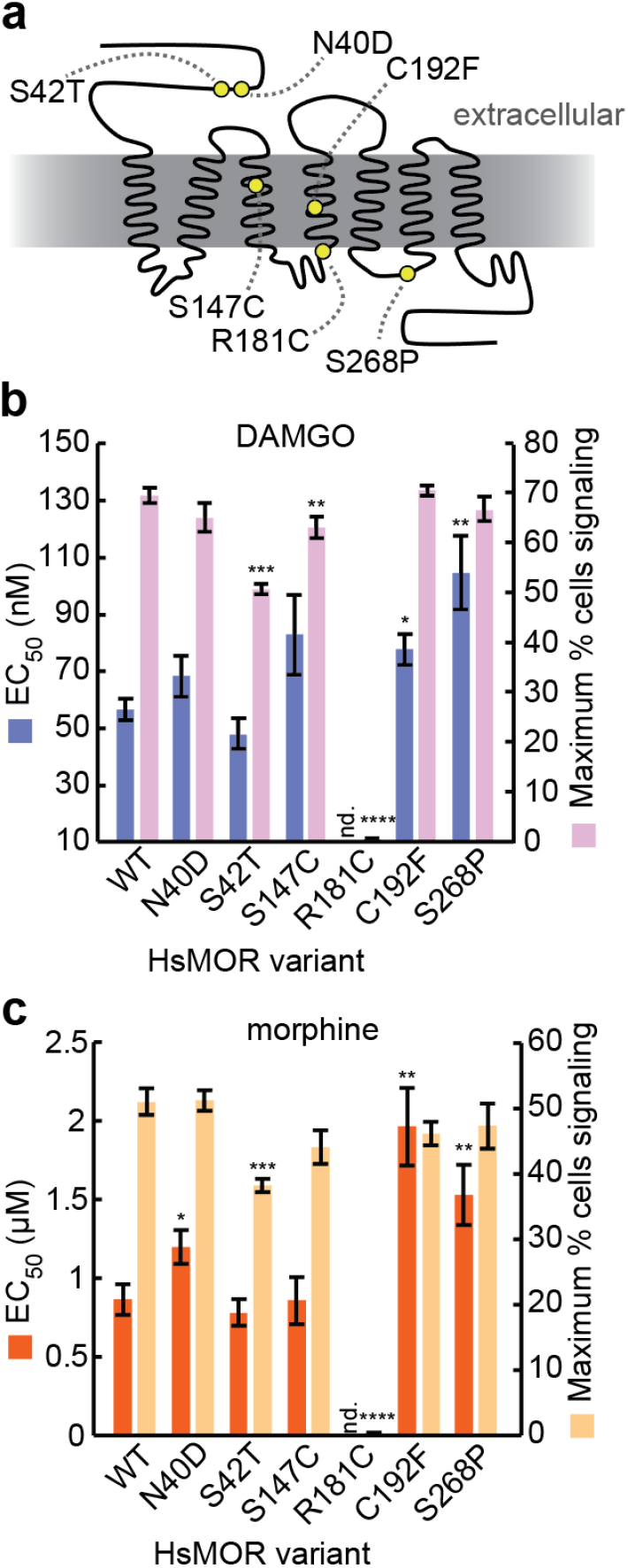
The effects of HsMOR missense mutations on opioid biosensors activity. **(a)** HsMOR snake plot with previously identified missense mutations. Biosensors based on these mutants were assayed for DAMGO response **(b)**, or morphine response **(c)**, by flow cytometry. Paired one-way ANOVAs of DAMGO *pEC_50_*s and maximum percentage of cells signaling: n=6, >2956 cells/condition/replicate; P = 0.0002 and P < 0.0001 respectively; Dunnett’s tests against WT shown. Paired one-way ANOVAs of morphine *pEC_50_*s and maximum percentage of cells signaling: n=6, >4761 cells/condition/replicate; P = 0.0033 and P < 0.0001 respectively; Dunnett’s tests against WT shown. Error bars indicate SEM. *, P < 0.05; **, P < 0.01; ***, P < 0.001; ****, P < 0.0001.

Variant HsMORs were introduced into the biosensor background and response to DAMGO and morphine was measured (Figure 4B/C). In agreement with previous work, the signal transduction-defective HsMOR(R181C) mutant was unable to respond to either DAMGO or morphine^50–52^. While dramatic defects were clearly detected, alleles associated with subtle defects were also explored. Previous descriptions of the relatively common (8-16% frequency) HsMOR(N40D) allele are more ambiguous, alternately describing no effect on agonist affinities or decreased β-endorphin affinity, while decreased analgesic response to morphine has also been reported^50,53^. In our biosensor, the N40D variant did not differ in DAMGO response though it displayed a decrease in morphine sensitivity (EC_50_ +38%), consistent with the reported decrease in morphine-based analgesia. Another variant, S268P, has a disrupted phosphorylation site and has been associated with reduced G protein coupling and reduced internalization and desensitization. A HsMOR(S268P)-based biosensor displayed decreased sensitivity to DAMGO (EC_50_ +84%) and morphine (EC_50_ +75%), consistent with diminished G protein coupling and raising the possibility of native yeast kinases acting on exogenous GPCRs.

Ravindranathan *et al*. characterized other HsMOR variants that resulted in mild decreases (S42T, C192F) or an increase (S147C) in sensitivity to DAMGO and morphine^52^. Correspondingly, a HsMOR(S42T)-based biosensor displayed decreased signaling populations with both agonists, and a HsMOR(C192F)-based biosensor had significantly lower sensitivities to DAMGO (EC_50_ +37%) and morphine (EC_50_ +125%). However, HsMOR(S147C) did not show improved sensitivity, instead resulting in a mild 10% decrease in the DAMGO-induced signalling population. Thus, the HsMOR biosensor provides a powerful platform to screen variants for changes in activity, which could inform how patients will respond to opioid-based analgesics.

### Exploring the functional significance of HsMOR terminal domains

We further applied our platform to explore how additional HsMOR structural variants affect receptor activity and localization in yeast. Opioid receptor terminal domains are moderately conserved, often containing trafficking motifs, glycosylation sites and phosphorylation sites, collectively contributing to folding, localization and modification of activity^32,54^. We first made variants lacking putative trafficking motifs R_367_xR and L_389_xxLE, or all five putative N-linked glycosylation sites (Figure 5A). RxR motifs can bind the coatomer protein I (COPI) complex and have been shown to mediate delta opioid receptor ER/Golgi retention^55^, while LxxLE can be recognized by COPII, facilitating ER exit^56^. N-glycosylation aids in protein quality control and contributes to DOR and KOR folding, stability and trafficking^57–59^. In response to DAMGO and morphine, biosensors based on all variants displayed subtle decreases in sensitivity (1.6-2.6-fold), suggesting these regions do not greatly contribute to folding or trafficking of HsMOR in yeast (Figure 5B). Consistently, isoform 2 of HsMOR, which contains a LENLEAETAPLP>VRSL C-terminal substitution and therefore lacks the LxxLE motif, has a similar signaling profile to isoform 1 (Figure 2B). However, removal of the RxR motif and the N-glycosylation sites did decrease the percent of cells signaling by up to 40% and 28% respectively, highlighting their contribution to achieving optimal activity (Figure 5C). In line with the overall mild defects, GFP-tagged variants displayed wild type localization (Figure 5D).

**Figure 5.**
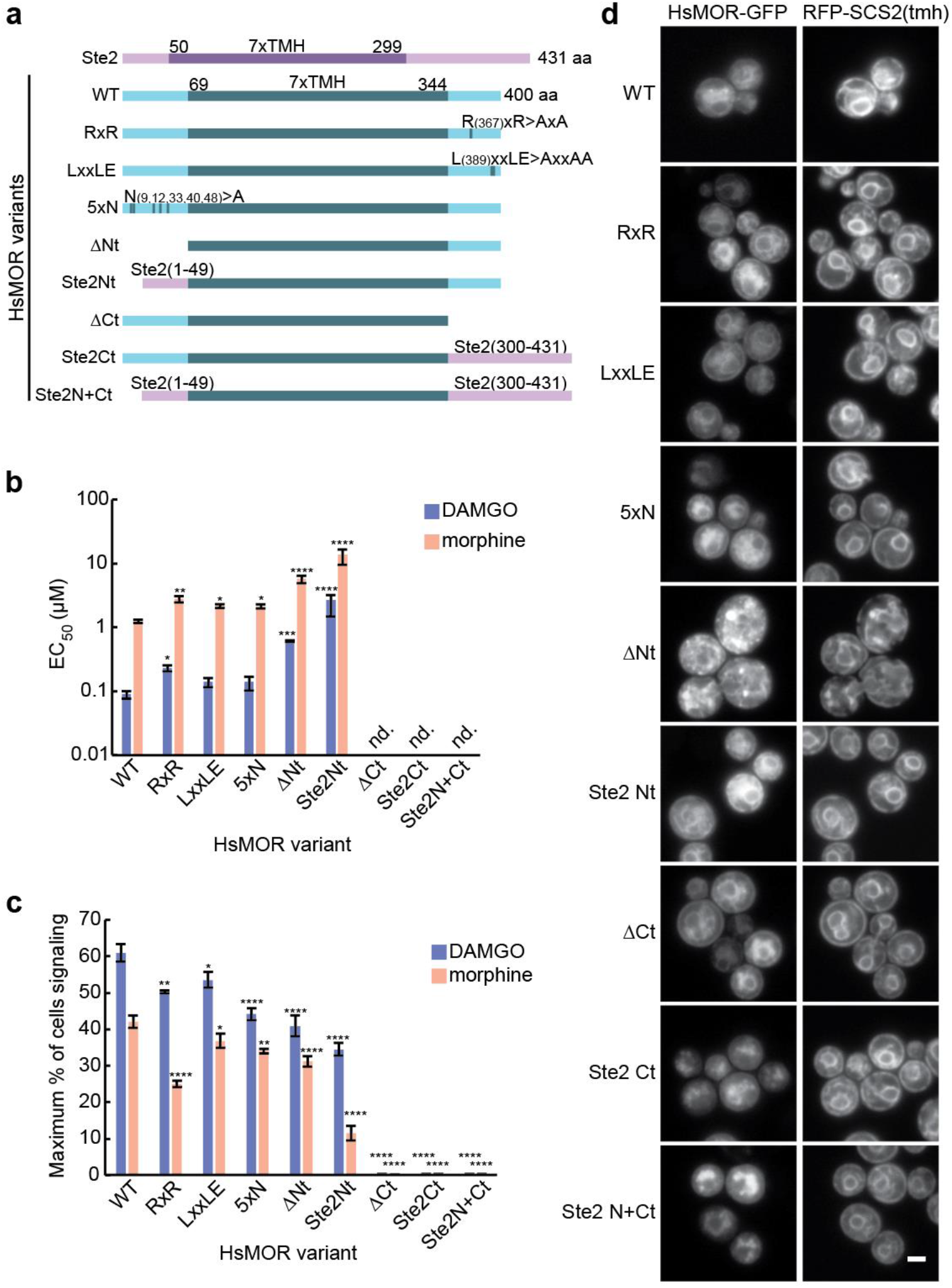
The functional requirements of HsMOR N- and C-terminal domains on biosensor activity. **(a)** Domains of Ste2, HsMOR and HsMOR variants with the conserved seven transmembrane helix (7xTMH) domains indicated. **(b)** DAMGO and morphine sensitivity of biosensors based on the indicated HsMOR variants. Unpaired one-way ANOVA of *pEC_50_*s: n=3, >7464 cells/strain/replicate; P < 0.0001 for both agonists; Dunnett’s tests against WT shown. **(c)** The effect of N- and C-terminal HsMOR mutations on the population of signaling biosensor cells. Unpaired one-way ANOVA: n=3, >7464 cells/strain/replicate; P < 0.0001 for both agonists; Dunnett’s tests against WT shown. **(d)** Imaging of C-terminally GFP-tagged HsMOR variants in a cholesterol-producing background with ER marker RFP-SCS2(tmh). Error bars indicate SEM. Scale bar is 2 μm. *, P < 0.05; **, P < 0.01; ***, P < 0.001; ****, P < 0.0001.

Next we tested complete removal of the HsMOR N- and C-terminal domains as well as substitution of these domains with those of the endogenous GPCR Ste2, as a small Ste2 N-terminal swap previously improved exogenous GPCR activity^6^. N-terminal deletion decreased DAMGO and morphine sensitivity by 6.8- and 4.6-fold respectively, in line with a previous report of a similar deletion causing a 3.3-fold drop in DAMGO affinity in HEK 293 cells^60^ (Figure 5B). Thus, the moderate functional contribution of the N-terminus appears conserved. In contrast with previous Ste2 swaps, complete substitution of the HsMOR N-terminus with that of Ste2 also decreased receptor function, reducing HsMOR DAMGO sensitivity 30-fold and decreasing the morphine signaling population by 72% (Figure 5B/C). However, unlike the N-terminal deletion, which displayed aberrant localization to ER-associated puncta, the N-terminal substitution displayed a wild type localization (Figure 5D). This suggests the Ste2 N-terminus is sufficient for maintaining localization and that localization poorly correlates with function.

C-terminal domain deletions or Ste2 substitutions also displayed a disconnect between localization and function as they showed no activity while maintaining nearly wild type localization, though with increased vacuolar pools (Figure 5B/C/D). The failure of the C-terminal mutants to signal was unexpected as a similar C-terminal deletion displayed only a small reduction in DAMGO sensitivity when expressed in CHO cells^61^. While this may indicate more stringent requirements for activity in yeast, the C-terminal deletions used here disrupt the short cytosolic helix (helix 8) next to the transmembrane domain that, while not involved directly in G protein binding or signal transduction, may contribute to the functional conformation of the receptor^62,63^. Taken together our results show our biosensors can be used to assess how domains and motifs contribute to function, and highlight the difficulty in linking activity to localization.

### Modifying membrane sterols alters HsMOR biosensor function

Cholesterol biosynthetic intermediates are typically present in plasma membranes at low concentrations, and accumulations are linked to developmental and neurological defects^64^. Still, relative proportions of cholesterol and its biosynthetic intermediates can vary based on tissue^65^. It remains unclear to what extent these intermediates can fulfill the roles of cholesterol in promoting GPCR activity. Profiles of sterol intermediates may exist that further promote GPCR signaling in yeast without cholesterol-associated growth and transformation defects^22^.

To search for sterol profiles that could improve HsMOR-based biosensor performance, we attempted to humanize the cholesterol biosynthetic pathway by introducing genes DHCR24, EBP, SC5DL and DHCR7 in a reconstructed erg2/3/5/6Δ biosensor background using another type of GFP, ZsGreenl, as the reporter (Figure 1D). Initially the human genes were introduced at the ergosterol biosynthesis gene loci, and driven by the native yeast promoters (erg5Δ::Hs.DHCR24, erg6Δ::Hs.DHCR7, erg3Δ::Hs.SC5DL, erg2Δ::Hs.EBP). GC-MS analysis of sterols showed this humanized strain generated the intermediates zymosterol, dehydrolathosterol, and 7-dehydrodesmosterol, while the products of DHCR7 and DHCR24 activity failed to accumulate (Figure 6B/C; Supplementary Figure 4).

**Figure 6.**
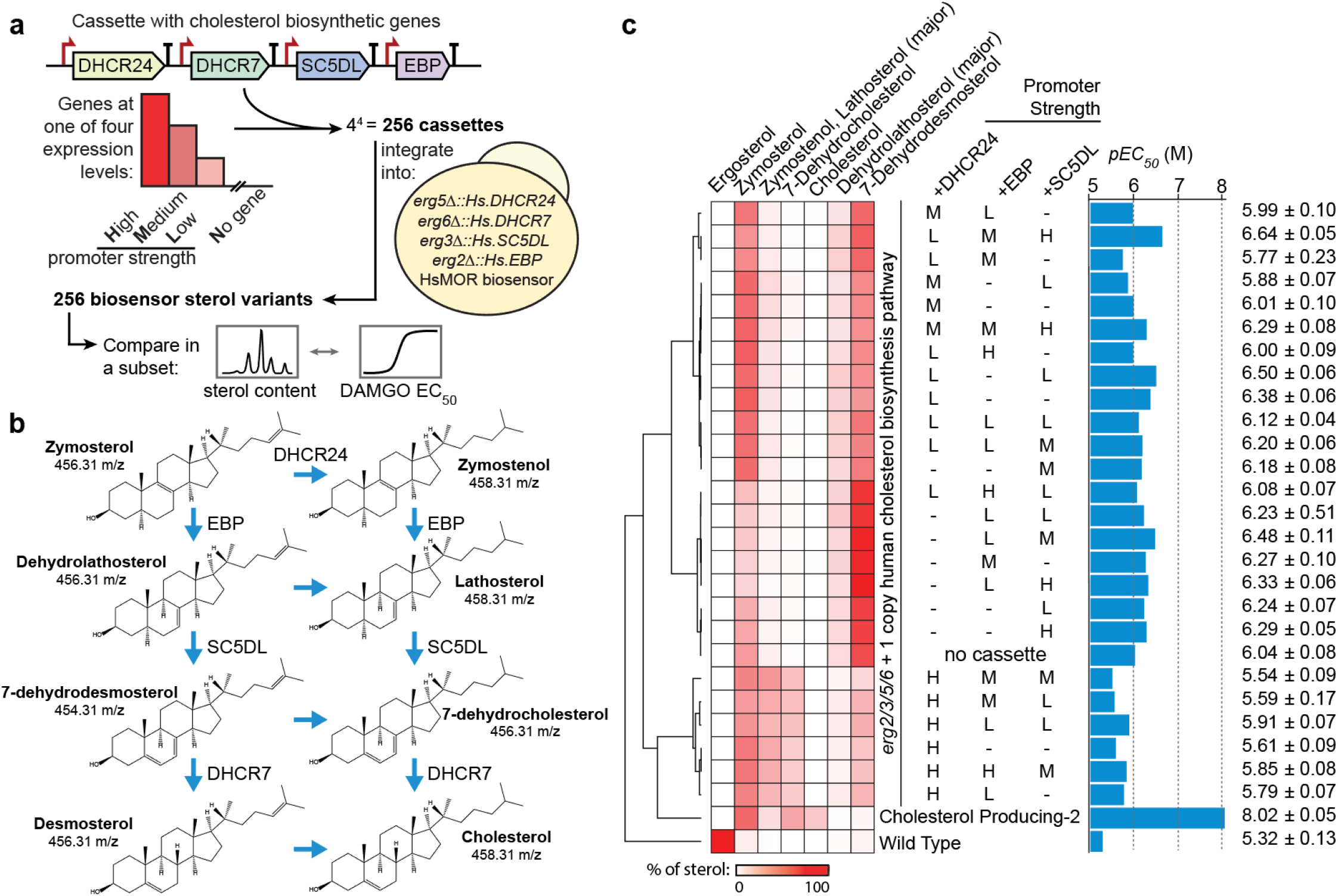
The effect of membrane sterol composition on opioid biosensor signaling efficiency. **(a)** Construction of an array of putative biosensor strains with systematically varied promoter strength for the cholesterol biosynthetic genes downstream of zymosterol. Cassettes containing these genes under different promoters were integrated into an HsMOR biosensor background with an additional copy of these enzymes in place of the final ergosterol biosynthetic genes (*ERG2/3/5/6*). **(b)** Cholesterol biosynthesis intermediates after zymosterol. **(c)** Sterol analysis and full dose responses with DAMGO of 39% of biosensors from the collapsed screen. Percentages of total sterol content for the intermediates were determined and *pEC_50_* concentrations were calculated.

Next we integrated in the yeast genome cassettes containing additional copies of the cholesterol biosynthetic genes under high, medium, or low strength yeast promoters to improve expression and generate strains with modified sterol profiles (Figure 6A). Of the 256 combinations, 249 were successfully constructed and assayed for response to 10 μM and 1 μM DAMGO, the concentrations roughly required to reach the E_max_ and EC_50_ in the wild type background (Figure 1F). Responses ranged from 21% - 61% and 8% - 47% of cells signaling at 10 μM and 1 μM DAMGO respectively. Human DHCR7 was found to be inactive, confirmed by failure of a Wilcoxon signed-rank test (P > 0.05) comparing the strong DHCR7 expression and no gene conditions, and the absence of products from DHCR7 activity in downstream sterol analyses (Supplementary Figure 3A, data not shown). Therefore, we excluded DHCR7 from our analysis and selected 39% of strains from this collapsed set for membrane sterol composition analysis (Supplementary Figure 3A). GC-MS analysis revealed that most variation was in 7-dehydrocholesterol, zymosterol, zymostenol, and lathosterol (Figure 6C). Subsequently, dose responses using the agonist DAMGO were performed in duplicate, and EC_50_ values for each strain were determined (Figure 6C). We also performed these analyses in similarly constructed biosensors with a wild type ergosterol-producing background and an alternative cholesterol-producing (erg5Δ::TDH3pr-Dr.DHCR7 erg6Δ::CCW12pr-Dr.DHCR24; Cholesterol Producing-2) background.

Hierarchical clustering identified trends in the composition of sterol intermediates. In particular, variations in DHCR24 promoter strength led to the largest changes in sterol composition, with higher promoter strength correlating with decreased HsMOR sensitivity (Figure 6C). The single copy of DHCR24 in the base strain proved insufficient to produce zymostenol, lathosterol, and 7-dehydrocholesterol. Accordingly, the presence of these intermediates correlate with higher EC_50_ values. A linear regression analysis on the sterol intermediate percentages and EC_50_s reinforced the relationship between sterol composition and signaling, finding a strong correlation (Supplementary Figure 3D). The cholesterol-producing biosensor strain proved disproportionally more sensitive with an EC_50_ approximately 24 times lower than the most sensitive strain identified from the screen (Figure 6C).

### Sterol modifications improve human class A GPCR function in yeast

To explore how broadly cholesterol improves functional expression of human GPCRs in yeast, we introduced seven different GPCRs into wild type, cholesterol-producing and sterol intermediate biosensor backgrounds. These receptors belong to three GPCR classes, all can couple with the Gvo chimera, and four of them, HTR4B, GLP1R, SSTR5, and FPR1, have been shown to function in yeast^66–69^ (Figure 7A). Of the resulting putative biosensors, all strains with class A receptors showed response to their cognate agonists at 10 μM and lower, whereas no class B or C receptors signaled in any sterol background (Figure 7C). Of the receptors reported to be active in yeast, only GLP1R failed to signal, possibly due to the use of different assays.

**Figure 7.**
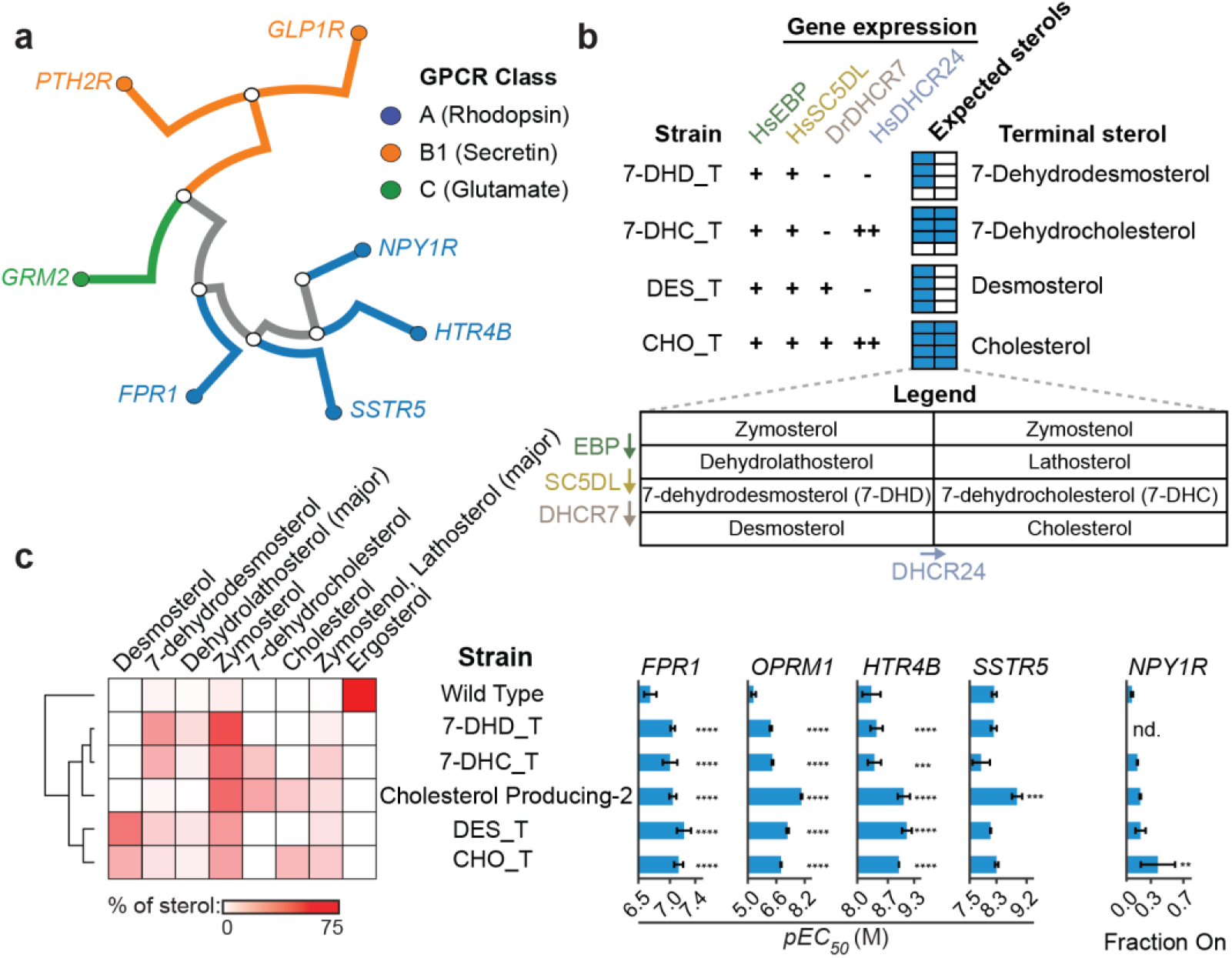
Effect of cholesterol on the activity of different human GPCRs. **(a)** Phylogenetic relationships of GPCRs tested. **(b)** Strains, designated by their terminal sterol, were built to produce all sterols up to the indicated terminal sterols by introducing sterol biosynthetic genes under native ergosterol promoters (+) or strong yeast promoters (++) in a *erg2/3/5/6* biosensor background. (-) indicates no gene is present. The expected sterol space is diagrammed in the expected sterols legend. **(c)** Sterol profiles of strains with the indicated sterol backgrounds were determined. Biosensors based on expressing *HTR4B, FPR1, OPRM1, SSTR5* and *NPY1R* in all sterol backgrounds were assayed in the presence of their agonists by flow cytometry to construct dose-response curves used to determine pEC_50_s and the maximum fraction on. Where responses were too weak to determine pEC_50_, fraction on is reported. Unpaired one-way ANOVA: n=3, >10,000 cells/strain/replicate; P < 2.2e-16 for *HTR4B, FPR1, OPRM1, SSTR5* assays, P = 0.02518 for *NPY1R* assay. Dunnett’s tests against WT shown. *, P < 0.05; **, P < 0.01; ***, P < 0.001; ****, P < 0.0001. *OPRM1*, Wild Type and *OPRM1*, Cholesterol Producing-2 data are replotted from Figure 6C.

In order to more generally test these GPCRs’ activities with respect to membrane sterols, as our promoter screen only sampled a subset of intermediate sterols (Figure 6C), we engineered strains targeting specific terminal sterols (Figure 7B). This was achieved by selectively expressing a subset of the cholesterol biosynthetic genes in an *erg2/3/5/6* background. Using this panel of strains, we measured dose-response curves for the active receptors (FPR1, SSTR5, HTR4B, HsMOR) in each sterol background; a dose response could not be measured NPY1R since it only responded to neuropeptide Y concentrations approaching 10 μM. Remarkably, sensitivities of all biosensors were greater in one or more of the modified sterol strains in comparison to the ergosterol-producing wild type strain (Figure 7C). Increases in pEC_50_s with the non-opioid receptors ranged from 2.8-fold (FPR1) to 6.5-fold (5-HTR4) in the best performing strains. For NPY1R, the maximum fraction of cells signaling increased 5.7-fold in the top-performing strain compared to the wild type strain. Thus, the activity of class A GPCRs in yeast can generally be improved by replacing ergosterol with cholesterol and related human sterol intermediates, with different specific sterols preferred by different receptors.

## DISCUSSION

We have shown that engineering yeast to produce cholesterol and other human sterols is an effective strategy for improving vertebrate GPCR activity in yeast, thereby enabling the generation of opioid biosensors with nanomolar sensitivities and expected agonist selectivities. This allowed us to evaluate the structural requirements for HsMOR function in yeast and recapitulate many defects associated with clinically relevant missense mutations. Systematic modification of the sterol biosynthetic pathway revealed that while the presence of upstream cholesterol intermediates can improve activity, a cholesterol producing background is most effective for HsMOR function. The presence of cholesterol and related human sterol intermediates also improved the function of several other GPCRs (FPR1, HTR4B, SSTR5 and NPY1R) indicating that modification of sterols is a general tool for the functional expression of animal GPCRs in yeast.

GPCRs can require cholesterol for normal function or regulation, likely due to both specific GPCR-cholesterol interactions^20^ and non-specific effects such as increased membrane fluidity or the facilitation of lipid subdomains^70^. By comparing GPCR activity in cholesterol- and ergosterol-producing yeast we indirectly assessed the extent to which cholesterol is specifically required for human GPCR activity. Remarkably, cholesterol increased the sensitivity of all tested GPCRs, even though only HsMOR has reported cholesterol-dependence. This suggests that cholesterol often improves human GPCR function beyond a non-specific requirement for sterols in the membrane. Conversely, non-native sterols may actively disrupt function, as Lagane et al. could detect DAMGO binding by HsMOR in yeast lysates only after ergosterol depletion with methyl-β-cyclodextrin^9^. This effect likely contributed to the performance improvements of biosensors producing sterol intermediates. Taken together, the frequency with which GPCRs have evolved to utilize direct interactions with native sterols may be underestimated.

Though many GPCRs benefited from the presence of cholesterol, HsMOR displayed the greatest improvements in sensitivity (Figures 1 and 7). HsMOR cholesterol dependence was expected as cholesterol is bound in HsMOR crystal structures^62^ and there is evidence that cholesterol directly promotes an active conformation^71^, partitions the receptor into more functional subdomains^72^, and aids in dimerization^73^. Though there is evidence that this receptor could be directly interacting with cholesterol in yeast, the degree to which this is occurring and the mechanism by which this improves activity remains to be resolved. The milder cholesterol dependence of the other receptors likely reflects a more limited potential for cholesterol binding. While cholesterol often improves activity, it has been shown to disrupt the activity of some receptors including the M2 muscarinic acetylcholine receptor^75^, type 1 cannabinoid receptors^76^, and rhodopsin^77^. It remains to be determined if the activity of these receptors is similarly disrupted in a cholesterol-producing yeast strain and if rules can be developed to predict which receptors will most benefit from conversion of ergosterol to cholesterol.

Small differences in sterol structure appear to have significant effects on HsMOR signaling. Screening HsMOR-based biosensors producing different sterol intermediates revealed that any combination of cholesterol intermediates increases sensitivity relative to ergosterol, though lathosterol and zymostenol were least beneficial. This was surprising given that in humans enrichment of these intermediates, including zymosterol, lathosterol and 7-dehydrocholesterol, is disruptive and linked to several diseases^64^. Indeed, one GPCR, HTR1A, can be disrupted by increasing 7-dehydrocholesterol levels to mimic Smith-Lemli-Opitz Syndrome^78^. Our sterol intermediate biosensors offer surrogate strategies to screen for other similarly disrupted GPCRs.

Our cholesterol-rich background enabled all opioid receptors to signal, generally with expected agonist specificities, allowing us to establish interspecies conservation of receptor function. The mammalian opioid receptors consistently displayed specificities similar to those of humans, whereas the responses of less related receptors were more variable. Some receptors such as the flying bat MOR, python DOR, or the zebrafish KOR only weakly responded to some of the agonists. In contrast, the bearded dragon KOR had the strongest response to KOR agonists and was also able to respond to the MOR-specific agonist DAMGO. The zebrafish DOR and KOR-based biosensors each showed no response to one of the two type-specific agonists tested, in line with previous work indicating the zebrafish DOR responds more strongly to general agonists than MOR, KOR, or DOR-specific agonists^79^. Indeed, a previous model suggests that there is an increased rate of divergence of mammalian opioid receptors from ancestral receptors relative to those of fish and reptiles, leading to more robust agonist specificities^32^. While our data partially supports this model, we find agonist specificities to be widely conserved.

While specificity was well conserved, opioid biosensors were on average 54-fold less sensitive than values previously determined for receptors in more native environments (Figure 2C). Only the HsDOR met-enkephalin response outperformed reported sensitivities with an EC_50_ of 10nM, a 3-fold improvement. Notably, the drop in sensitivity of MORs in our biosensors was largest and roughly ten times greater than that of DORs, a substantial difference given their close evolutionary and structural relationships (Figure 2A). Perhaps MORs, which appear to have the highest evolutionary rate^32^, diverged to require additional features of the vertebrate environment for full function. Species of origin was poorly correlated with sensitivity as the origins of the most sensitive mu, delta, and kappa receptors were diverse: mice, humans, and bearded dragons respectively. This indicates that opioid receptor sensitivity may be heavily influenced by sporadic mutations that coincidentally improve performance in yeast.

Thus, there is room to improve opioid biosensor performance, perhaps by further adjusting the biosensor environment or its components. Here, GPCRs were generally codon optimized to improve yeast expression. However, additional tests on a subset of six opioid biosensors, two of each receptor type, found that native genes improve sensitivity by as much as 31-fold for the SNC80 response of PbDOR, and 3.9-fold on average (Supplementary Figure 1). Though the sensitivity improvement was tempered by a 1.4-fold reduction in percent of cells signaling, using GPCRs with native codons may be beneficial overall. This may be because native genes contain rare codons, which could decrease the rate of translation, potentially promoting opioid receptors achieving optimal folds. Other approaches to improve biosensor activity may include strengthening the link to the pheromone response pathway, adding potential chaperones, or performing unbiased screens for yeast deletions that improve activity. Introducing enzymes responsible for post-translational modifications such as palmitoylation^54^ or attempting to adjust yeast membrane thickness^80^ may also be helpful. Alternatively, applying slower biosensor assays that allow greater signal accumulation, such as the 24 h β-galactosidase method used by Olesnicky *et al*., could improve sensitivity^25^.

Our opioid biosensors and sterol-modified biosensor backgrounds have many applications. The speed and low cost of using our opioid biosensors for screening compounds for receptor type-specific activation should make them an attractive tool to bridge computational docking studies^81^ and more costly screens in human cell lines based on protein complementation^40^ or bioluminescence resonance energy transfer^82^. Currently our opioid biosensors are unable to measure modes of signaling beyond G protein activation, such as β-arrestin recruitment, which is thought to cause many of the side effects of opioids. This makes the biosensors less useful for drug discovery efforts which are focused on identifying compounds that display biased agonism towards G protein activation. However, our biosensors are compatible with the PRESTO-Tango^83^ system for detecting GPCR-β-arrestin interactions, which would allow future biosensors to detect biased agonism. By increasing throughput of production assays from hundreds to thousands, these biosensors will also aid in the ongoing development of opiate production strains^84^. Furthermore, it may be possible to adapt the opioid biosensors to field tests for opioid detection. Colorimetric assays based on yeast biosensors have been reported previously^85^, and in principle our biosensors could be used to test a sample for the degree of opioid activity independent of identifying the compounds present. This may enable testing kits that could be used to assess the amount of a sample likely to cause an overdose. Beyond opioid biosensors, our sterol modified platform should enable the expression of many other human GPCRs in yeast, generating an array of new biosensors and tools for the deorphanization of GPCRs.

## MATERIALS AND METHODS

### Strains and Plasmids

Strains and plasmids are listed in Supplementary Table 1 and 2. Strains were derived from BY4741^86^ using CRISPR-Cas9 as follows. A Cas9 (CEN6 *URA3*) vector was constructed using components of the Yeast Toolkit^87^, with *pPGK1*-Cas9-*tENO2* and up to four sgRNAs expressed from a tRNA^phe^ promoter with a 5’ HDV ribozyme site and a *SNR52* terminator. Alternatively, a Cas9 vector derived from the vector described in Ryan *et al*. was used^88^. Strains were constructed by transforming yeast with a Cas9 vector, unique protospacers guiding Cas9, and a double stranded repair template introducing deletions or modifications. Deletions and modifications were confirmed by colony PCR and sequencing respectively. All protospacers and repair template sequences are listed in Supplementary Table 3. Yeast were transformed using either the Zymo Research EZ Yeast Transformation II Kit (cat. T2001) or a modified Gietz protocol^89^.

Plasmids were constructed using Golden Gate assembly of components from the Yeast Toolkit^87^ and elsewhere. Opioid receptors were all expressed from the same 2μ HIS3 backbone assembly (ConLS’-*CCW12p*-<u>GPCR</u>-*SSAlt*-ConRE’-*HIS3-2μ-KanR*-ColE1) while *FPR1* was on a similar vector with a *TDH1* terminator and other GPCRs were expressed from a ConLS’-*CCW12p*-<u>GPCR</u>-*SSAlt*-ConRE’-URA3-2μ-KanR-pI5a backbone. GPCRs were ordered as either gblocks from IDT or clonal genes from Twist Biosciences. GPCR sequences are listed in Supplementary Table 4 and were yeast codon optimized unless specified as non-Codon Optimized (nCO).

### Sterol extraction

Yeast strains were grown to either mid-log (8hrs) or saturation (48 hrs) from single colonies. Since the growth rates of these strains were different, wet weights were adjusted to 50mg and 150mg for the 8hr and 48hr timepoints respectively. These were then suspended in glass tubes containing 3 ml of 10% w/v methanolic KOH. The tubes were flushed with nitrogen gas and capped before incubating at 70°C for 90 min. Samples were cooled to room temperature before 1 ml of water and 2 ml of n-hexane were added and vortexed. The hexane phase was transferred to glass vials and the extraction process was repeated. Combined extracts were dried under nitrogen and derivatized by adding 50μl N,O-Bis(trimethylsilyl)trifluoroacetamide:Trimethylchlorosilane (BSTFA, 1% TMCS) and incubating at 60°C for 30 min. Derivatized samples were dried under nitrogen or by vacuum centrifugation for −30 min, and finally suspended in ethyl acetate for GC-MS analysis.

### GC-MS analysis of sterols

Derivatized sterol extracts and standards were analyzed on an Agilent Technologies 5977 GC/MSD equipped with a Agilent J&W DB-1MS UI capillary column with 45m in length, 0.25 mm inner diameter and 0.25 μm phase thickness (phase-100% dimethylpolysiloxane). Sterols from 1 μl injections were separated using an initial oven temperature of 40°C for 1 min followed by a 20°C/min ramp to 320°C, which was held for 12 min (constant helium flow of 1 ml/min). The mass spectrometer source and transfer line temperatures were set at 260°C and 280°C, respectively and the GC inlet was operated in splitless mode. Mass spectral data was analyzed using MassHunter Workstation Software (Agilent). Parent and fragment ion counts were extracted at 129.3, 454.3, 456.3, 458.3, and 468.3 m/z using a window of +/- 0.5 m/z for analysis. Extracted Ion Chromatograms (EICs) were aligned, then individual sterols quantified as baseline-corrected peak areas across appropriate retention time windows for the following ions: 454.3, 7-dehydrodesmosterol; 456.3, 7-dehydrocholesterol, zymosterol, 7-dehydrolathosterol; 458.3, cholesterol, zymostenol+lathosterol; 468.3, ergosterol. Relative sterol abundances were calculated as the percentage of total ions detected for the set of measured sterols. Ambiguities between 7-dehydrocholesterol and desmosterol were resolved by examination of the 129/456 fragment ion ratio, and assignments confirmed using purified standards as shown in Supplementary Figure 4.

### Plate reader signaling assay

Yeast were grown overnight in synthetic selective media and back-diluted 1:10 into media, with agonists as indicated, in Falcon 96 well micro titer plates to 100 μL final volumes. Cells were shaken at 30°C for either 3 hours (alpha mating factor tests) or 8 hours (DAMGO tests) prior to measurement on a CLARIOstar plate reader (BMG Labtech). Values for OD600 and green fluorescence (excitation 469 nm ±13 nm, emission 508 nm ±15 nm) or red fluorescence (excitation 527 nm ± 27 nm, emission 622 nm ± 30 nm) were collected for each sample.

### Flow cytometer signaling assay

Overnight cultures grown in synthetic selective media were back-diluted 1:10 into fresh media containing the agonist being tested to a final volume of 100 μL in a Falcon 96 well microtiter plate. Cells were shaken at 300 rpm for 8 hours (or 6 hours for alpha mating factor tests) prior to measurement on an BD Accuri C6 flow cytometer. Either 10000 events, or those within 15 μL of the culture, were recorded. For alpha mating factor response measurements the mean green fluorescence of the complete, ungated population was determined and used to calculate fold induction of fluorescence. Otherwise, within an experiment the biosensor that was brightest in its inactive state (no agonist) was used to establish an arbitrary green fluorescence intensity threshold such that 0.1-1% of cells were brighter than the threshold. This threshold was propagated to all conditions within the experiment and the percentage of the cells within each measurement that exceeded the threshold were recorded as the percentage of cells signaling. The percentage of cells signaling was exported to construct 4 parameter dose-response curves within Prism 8 (GraphPad) and calculate EC50s, IC50s and the maximum percentage of cells signaling within a biosensor-agonist condition.

Alternatively, for Figure 6, overnight cultures were back-diluted to an OD600 of approximately 0.2. The agonist was added upon dilution and cells were grown for 8 hours in 96-well deep well plates at a volume of 500 μl at 30°C with shaking at 1000 rpm. 10000 cells of each sample were analyzed using a Sony SP6800 Spectral Analyzer.

### Microscopy

Log phase yeast grown in synthetic selective media were mounted on slides and imaged using a DMi6000B microscope (Leica Microsystems) with an HCX PL APO 63x oil objective, an Orca R2 CCD camera (Hamamatsu) and Volocity software (PerkinElmer). Images were processed using Fiji^90^ and Photoshop CC (Adobe).

## ABBREVIATIONS

au: arbitrary units
αMF: alpha mating factor
DAMGO: [D-Ala^2^, N-MePhe^4^, Gly-ol]-enkephalin
DOR: delta opioid receptor
ER: endoplasmic reticulum
G protein: guanine nucleotide-binding protein
GPCR: G protein-coupled receptor
KOR: kappa opioid receptor
MOR: mu opioid receptor
PRP: mating pheromone response pathway
SD: standard deviation
SEM: standard error of the mean
tmh: transmembrane helix

## ACKNOWLEDGEMENTS

The authors thank Ian Riddington and the mass spectrometry facility at the University of Texas at Austin for their feedback and assistance with GC-MS sample preparation and analysis, and Josh Lutgens at the University of Texas at Austin for setting up and automating combinatorial DNA assembly using an Echo acoustic liquid handling robot. This study was financially supported by FRQNT Team and NSERC Discovery grants to V.J.J.M. and M.W.. B.D.M.B. was supported by a Concordia University Horizon Postdoctoral Fellowship, V.J.J.M. is supported by a Concordia University Senior Research Chair and M.W. is supported by a Canada Research Chair. Financial support was also provided by a Cooperative Agreement between the University of Texas at Austin and DEVCOM Army Research Laboratory to A.D.E., E.M.M., and J.D.G. (W911NF-17-2-0091). R.K.G was supported by the American Heart Association Predoctoral fellowship (#18PRE34060258). E.M.M. acknowledges additional support from the Welch Foundation (F-1515); Army Research Office (W911NF-12-1-0390); and NIH (R35 GM122480). The authors also acknowledge the Centre for Microscopy and Cellular Imaging funded by Concordia University, Montreal, Canada and the Canada Foundation for Innovation.

## CONTRIBUTIONS

B.D.M.B., C.J.M., J.D.G., M.W., and V.J.J.M. designed the research. B.D.M.B, C.J.M, R.K.G., D.R.B., O.R, W.C., B.M.F., E.C.G., and E.M.M. performed the experiments. V.J.J.M., M.W., A.D.E., E.M.M., and J.D.G. supervised the research. B.D.M.B. and C.J.M. wrote the manuscript with editing help from V.J.J.M., J.D.G., M.W., E.M.M. and R.K.G.

---

**Supplementary Figure 1.**
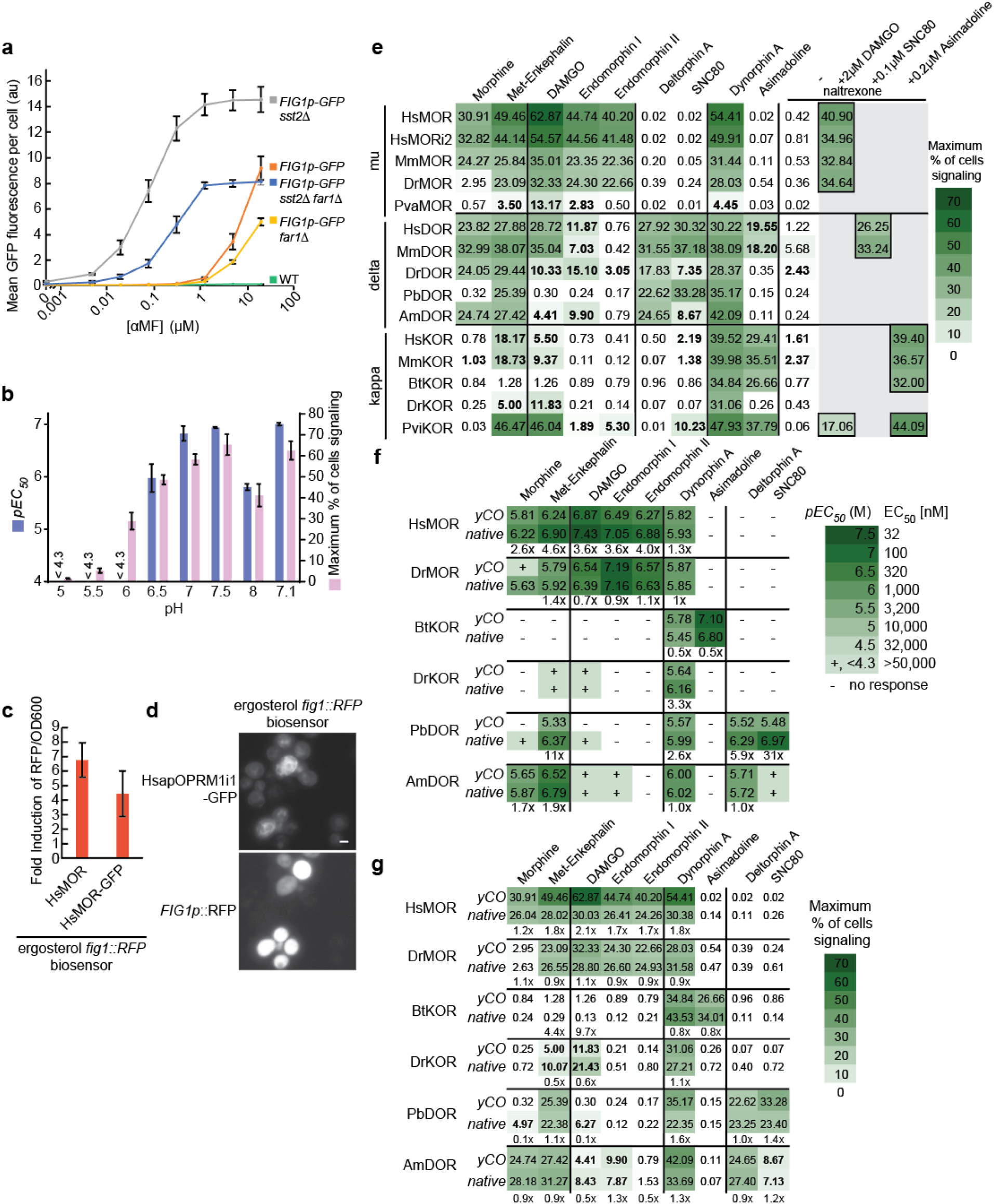
Additional metrics of biosensor activity. **(a)** Mean GFP fluorescence of strains treated with alpha mating factor for 6 h, measured by flow cytometry. n=3; 10,000 cells/strain/replicate. **(b)** pH dependence of the HsMOR-based cholesterol-producing biosensor response to DAMGO for 8 h measured by flow cytometry. Media buffered with 100 mM Tris. n=3, >8835 cells/condition/replicate. **(c)** The effect of GFP tagging the HsMOR C-terminus in an ergosterol biosensor background with a RFP reporter, as measured by a plate reader after 8h of 10 μM DAMGO agonist exposure. n=3. **(d)** Imaging of HsMOR-GFP biosensor after 6 h activation by 10 μM DAMGO. Scale bar is 2 μm. **(e)** Measurement of the maximum percent of biosensor cells signaling in the experiment presented in Figure 3B. n=3, >2073 cells/condition/replicate. **(f)** *pEC_50_* and **(g)** maximum percent of cells signaling for a subset of opioid biosensors based on yeast codon-optimized receptors compared with receptors expressed from genes with native codon usage. Measurements were taken after 8 h treatment with the indicated agonists, samples were measured by flow cytometry. Experiments run in parallel with that of panel **(e)**. n=3, >2073 cells/condition/replicate. Fold increase in native receptor EC_50_ and fold decrease in native receptor signaling populations are indicated. Error bars indicate SEM.

**Supplementary Figure 2.**
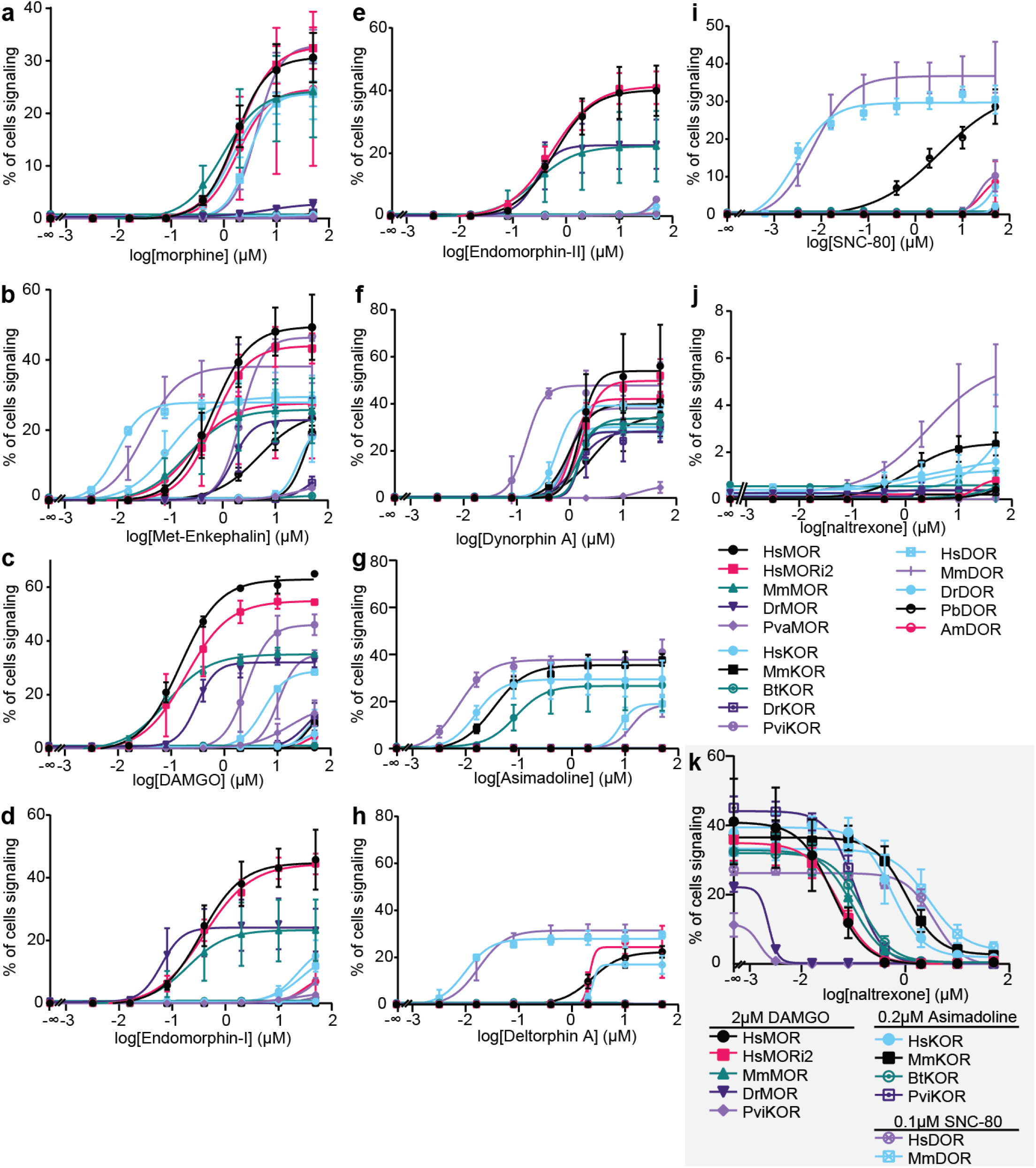
Dose-response curves of putative opioid biosensors against established agonists and antagonists. (**a-j)** Biosensor strain response to the indicated agonists were measured and fit using a 4 parameter nonlinear model. **(k)** The ability of naltrexone to block activation by the indicated agonists was determined. For **a-k**, n=3; >2073 cells/condition/replicate; error bars indicate SD.

**Supplementary Figure 3.**
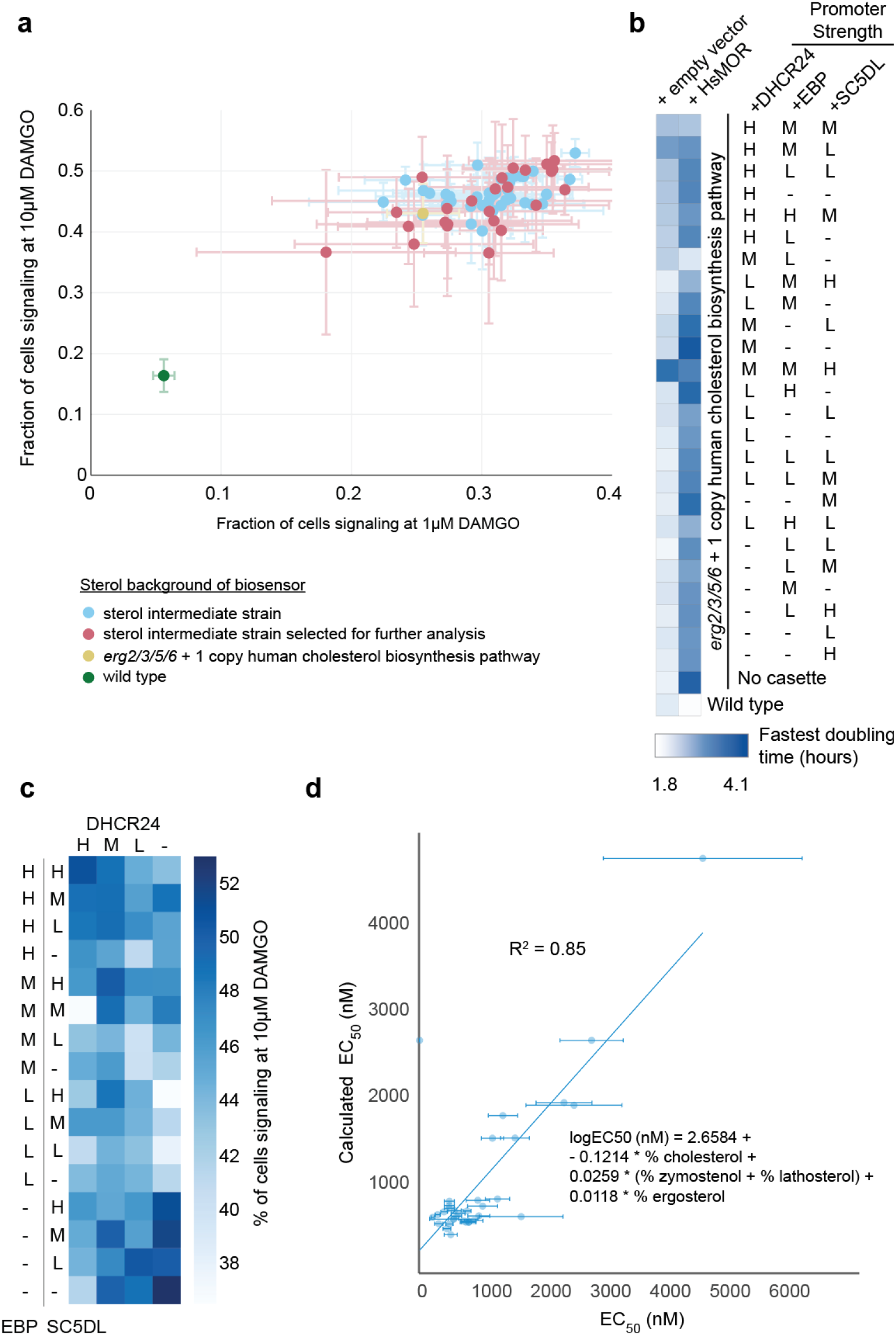
Characteristics of cholesterol biosynthesis intermediate screen strains. **(a)** Percent of cells signaling with 10 μM and 1 μM DAMGO. **(b)** Fastest doubling times of the strains chosen for further analysis, n=3. **(c)** The percent of cells signaling at 10 μM DAMGO in the sterol intermediate screen. Unpaired one-way ANOVA, P = 3.626e-14: n=4 or 3, 10000 cells/strain/replicate; P <1e-05 for all Dunnett’s tests against base strain. Results are averages across strains with varying (inactive) DHCR7 status. **(d)** The measured EC_50_s and sterol profiles of the shortlisted sterol intermediate strains were related using a linear regression. EC_50_s calculated based on sterol intermediate percentages with this regression were plotted against measured EC_50_s.

**Supplementary Figure 4.**
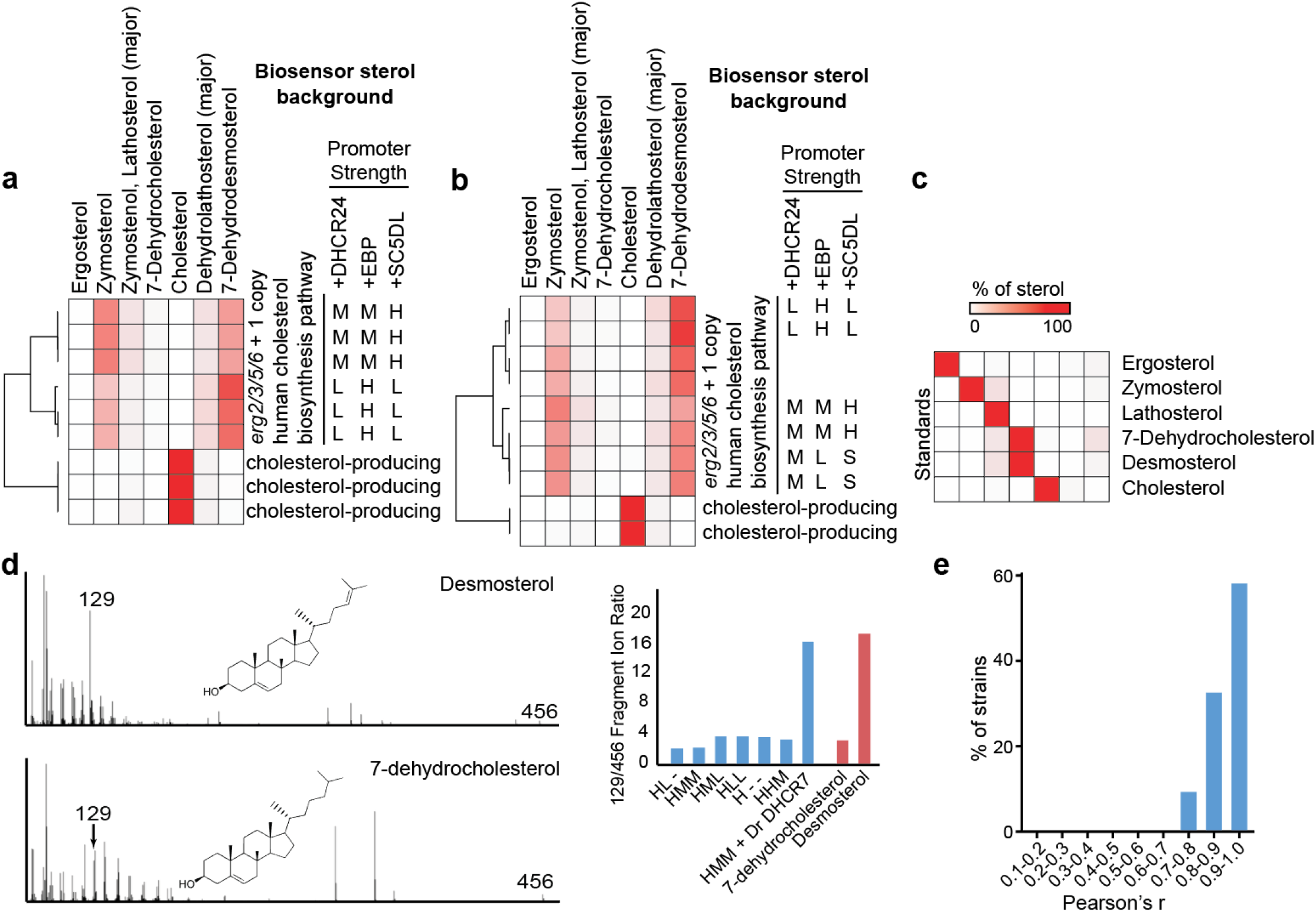
Consistency of sterol proportions across biological and technical replicates. Heatmaps representing percentages of total sterol content across **(a)** technical and **(b)** biological replicates. Percentage of total sterols were calculated as described in the Methods. Technical replicates constitute independent injections while biological replicates are samples from different colonies. **(c)** Heatmaps depicting percent of sterols present in standards as measured in **a** and **b**. **(d)** MS spectra for intermediate sterols with the same m/z ratio are distinguishable by characteristic peaks. (Left) Unlike desmosterol, 7-dehydrocholesterol lacks a strong peak at 129 m/z. (Right) Fragment ion ratios of 129 to 456 m/z across standards 7-dehydrocholesterol, desmosterol and engineered yeast strains. Low 129/456 ratios are indicative of 7-dehydrocholesterol while higher ratios correspond to desmosterol. Strains are all in the biosensor background with one copy of the human cholesterol biosynthetic pathway integrated. Names reflect promoter strengths for additional DHCR24, EBP, and SC5DL copies. **(e)** Distribution of Pearson r correlation coefficients between sterol composition measured from 43 strains grown either 8 h or 48 h.

